# Accelerated Antibody Discovery Targeting the SARS-CoV-2 Spike Protein for COVID-19 Therapeutic Potential

**DOI:** 10.1101/2021.05.31.446421

**Authors:** Tracey E. Mullen, Rashed Abdullah, Jacqueline Boucher, Anna Susi Brousseau, Narayan K. Dasuri, Noah T. Ditto, Andrew M. Doucette, Chloe Emery, Justin Gabriel, Brendan Greamo, Ketan S. Patil, Kelly Rothenberger, Justin Stolte, Colby A. Souders

**Author notes:** Corresponding authors &.

## Abstract

Rapid deployment of technologies capable of high-throughput and high-resolution screening is imperative for timely response to viral outbreaks. Risk mitigation in the form of leveraging multiple advanced technologies further increases the likelihood of identifying efficacious treatments in an aggressive timeline. In this study, we describe two parallel, yet distinct, *in vivo* approaches for accelerated discovery of antibodies targeting the SARS-CoV-2 spike protein. Working with human transgenic Alloy-GK mice, we detail a single B-cell discovery workflow to directly interrogate antibodies secreted from plasma cells for binding specificity and ACE2 receptor blocking activity. Additionally, we describe a concurrent accelerated hybridoma-based workflow utilizing a DiversimAb™ mouse model for increased diversity. The panel of antibodies isolated from both workflows revealed binding to distinct epitopes with both blocking and non-blocking profiles. Sequence analysis of the resulting lead candidates uncovered additional diversity with the opportunity for straightforward engineering and affinity maturation. By combining *in vivo* models with advanced integration of screening and selection platforms, lead antibody candidates can be sequenced and fully characterized within one to three months.

## Introduction

The global pandemic caused by severe acute respiratory syndrome coronavirus-2 (SARS-CoV-2), or coronavirus disease 2019 (COVID-19), has received unprecedented attention from the scientific community in an effort to rapidly develop efficacious treatments and vaccines. Within weeks of the emergence of viral pneumonia outbreaks in Wuhan, China, deep sequencing had identified the cause,^1^ and the resulting mobilization of widespread therapeutic and prophylactic discovery efforts ensued. The response to the COVID-19 pandemic mirrored that of other recent viral outbreaks, including, but not limted to, H1N1 influenza in 2009,^2^ Ebola Virus in 2014,^3,4^ and Zika Virus in 2015.^5^ Lessons learned from these public health threats helped guide the strategy for the accelerated response to COVID-19. In particular, the understanding that neutralizing antibody function is fundamental to combating disease progression^6^ helped streamline early antibody-based drug therapy discovery strategies.

Beyond direct therapeutic use, antibodies can help inform vaccine design to enable next-generation vaccine development with a focus on relevant viral epitopes.^7^ In general, the most valuable and broadly applicable antiviral antibodies are those that exhibit cross-reactivity to related viruses and are unaffected by escape mutant evolutionary pressures.^8^ These antibodies, which can function either alone or in combination with oligoclonal mixtures of non-competing antibodies,^9^ harbor basic properties like receptor blocking activity and high affinity. When taken in aggregate, these criteria are quite stringent and therefore necessitate efficient, high resolution screening strategies to identify valuable lead candidates.

This report highlights several different techniques and antibody discovery workflows leveraged in the discovery and characterization of antibody panels targeting the spike protein (S) of SARS-CoV-2. Across the different workflows, two separate mouse strains were immunized with the S1 subunit (which contains the receptor binding domain): a humanized strain to facilitate the discovery of fully human antibodies (Alloy GK mice), and an engineered mouse strain designed to elicit greater epitopic diversity and overall immune response (Abveris DiversimAb™ mice). Furthermore, two distinct upstream discovery methods were applied: a hybridoma discovery platform optimized for high-content screening and efficiency (Abveris Hybridoma Workflow), and a high-throughput state-of-the-art single B cell screening platform (Abveris Single B Cell Worflow enabled by the Berkeley Lights Beacon^®^). Final characterization and candidate analysis were performed on the Carterra LSA™.

## Methods and Results

Immunization of DiversimAb and Alloy GK mice was completed in 16 and 35 days, respectively; both protocols resulted in an appreciable immune response as indicated by detectable serum titer to the S and S1 proteins at serum dilution factors of at least 1:70,000 or higher (data not shown).

Following a high-efficiency electrofusion to generate hybridoma lines, resulting colonies were initially screened by ELISA to identify S protein-reactive clones. Preliminary clones of interest were subsequently screened via high-throughput biolayer interferometry (BLI) kinetic screening on the ForteBio Octet^®^ system to select candidates for scale up and antibody purification from hybridoma cultures. Simultaneously, sequencing of immunoglobulin genes was performed following a high-throughput hybridoma sequencing procedure and sequence analysis was performed using the Geneious Biologics software. Screening results from a subset of representative candidates are shown in Table 1.

**Table 1.**
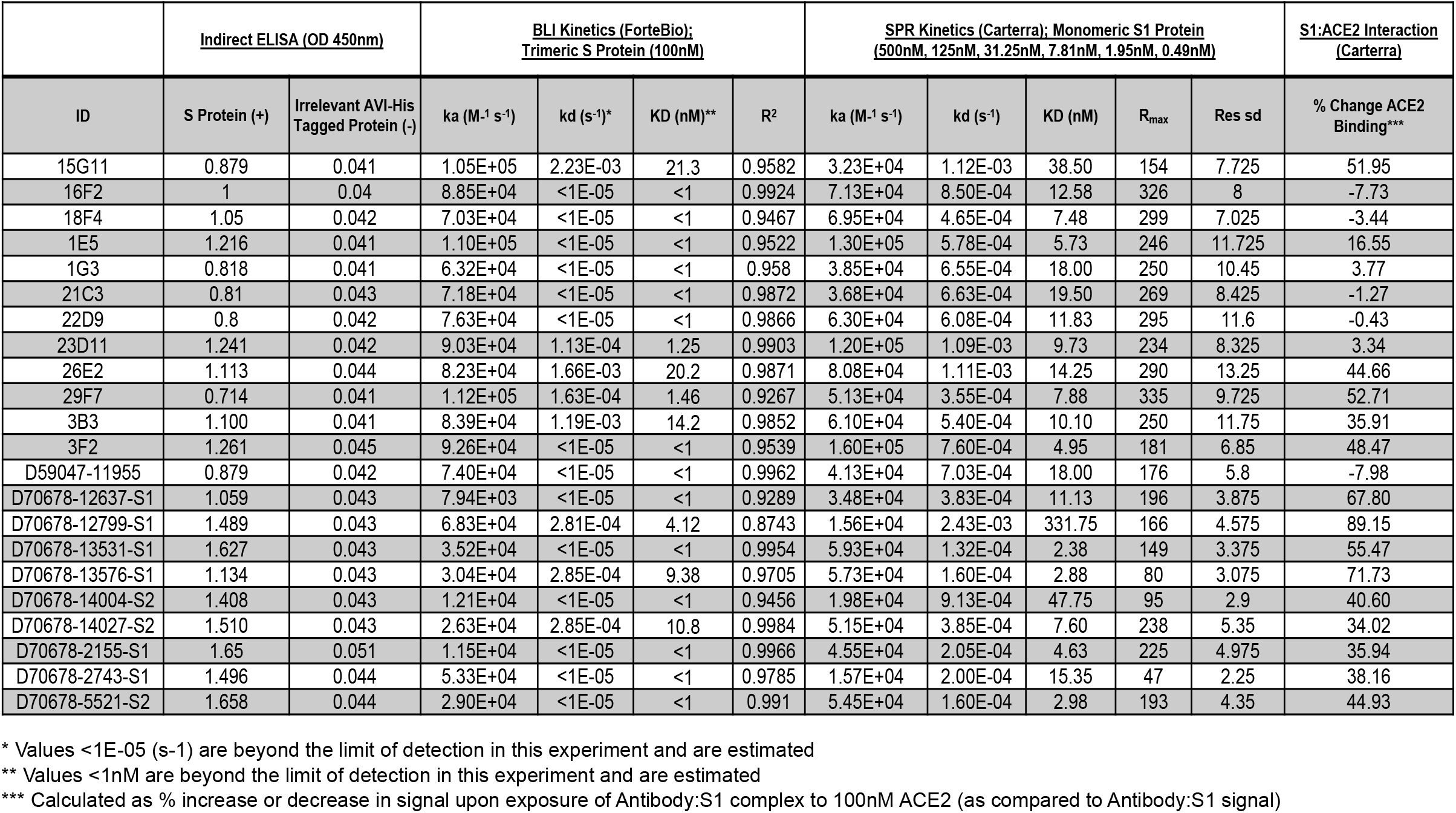
Binding characteristics of anti-S protein candidates, including ELISA binding, single point kinetic measurements to the trimeric S protein, affinity characterization to the monomeric S1 protein and effect on ACE2 binding to S1 protein (S1:ACE2 interaction blocking activity).

Concurrently, a single B cell discovery approach was employed whereby plasma cells were enriched from primary tissues prior to loading onto OptoSelect™ chips with the Berkeley Lights Beacon. Following single cell deposition into NanoPens, assay mixtures containing capture beads and fluorescently labeled target proteins were imported into the channels above the NanoPens. Throughout the course of the assay, antibody secreted from the plasma B cells diffused from the NanoPen chambers into the channel above. Upon bead binding, fluorescence from either directly labeled protein(s) or secondary detection antibodies was concentrated on the surface of the bead, resulting in the time-dependent development of fluorescent halos in the channels above the pens containing antigen-specific plasma cells (Figure 1).

**Figure 1.**
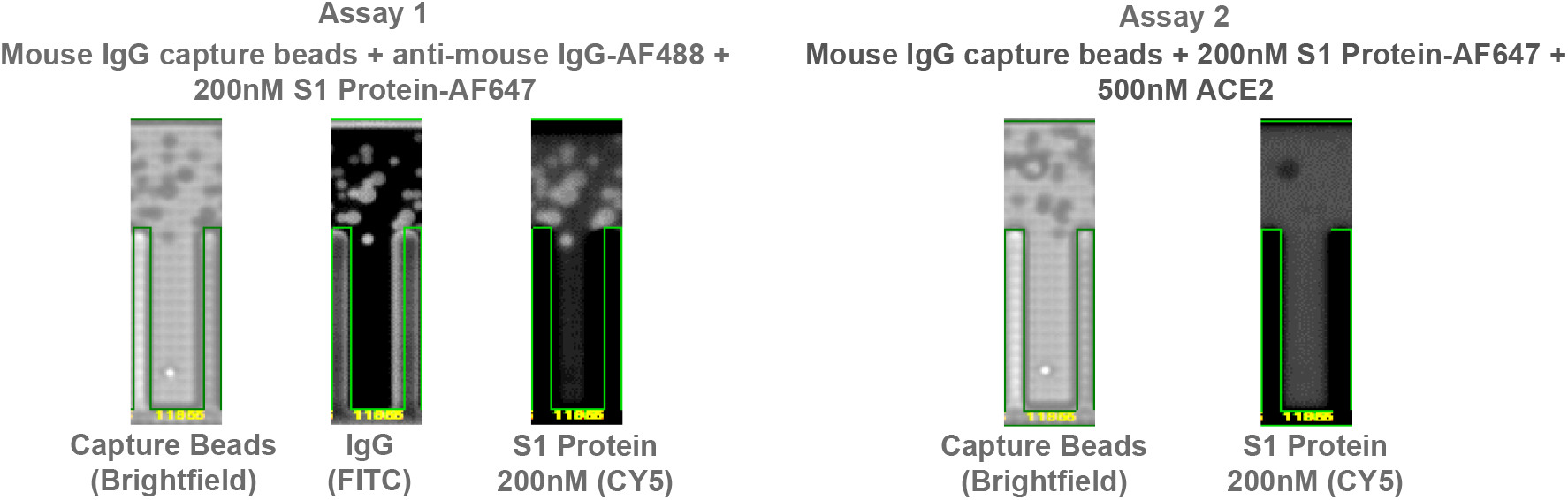
Example Beacon screening data for candidate D59047-11955. Anti-mouse IgG capture beads (brightfield image) were imported into the channel above the pens. Antibody secretion from a single B cell contained within a pen bound to capture beads at the mouth of the pen. In assay 1, antibody secretion was assessed by detection of total IgG in the FITC detection channel via binding of an anti-mouse IgG-AF488 conjugated secondary and simultaneously the specificity for S1 protein was determined in the CY5 detection channel with AF647 conjugated S1 protein at 200nM. In assay 2, binding competition between secreted antibody from the B cell and ACE2 receptor was assessed by precomplexing AF647-conjugated S1 protein with a molar excess of recombinant ACE2. A lack of antibody binding to S1 protein under these conditions demonstrated binding to a similar epitope as ACE2, thus indicative of a potential blocking candidate.

Plasma cells exhibiting on-Beacon binding profiles of interest were exported for immunoglobulin sequence capture and analysis with the Geneious Biologics software. The resulting naturally paired heavy and light chains were cloned into expression plasmids and recombinantly expressed. Purified antibodies were screened similarly to the strategy used for hybridoma candidates via ELISA and BLI with a subset of representative candidates displayed in Table 1.

High-throughput and high-content screening on lead candidates was performed on the Carterra LSA to elucidate kinetic profiles to the monovalent S1 protein (Table 1 and Figure 2). Full kinetic profiles were assessed in triplicate under regenerative and non-regenerative conditions with both purified antibodies and crude supernatant samples. Target S1 protein was used as an analyte in an ascending concentration series ranging from 0.49nM to 500nM with four-fold dilutions (Figure 2). All conditions (regeneration vs. non-regeneration and purified vs. crude antibody samples) yielded similar affinity values. The average resulting values from all asays are reported in Table 1.

**Figure 2.**
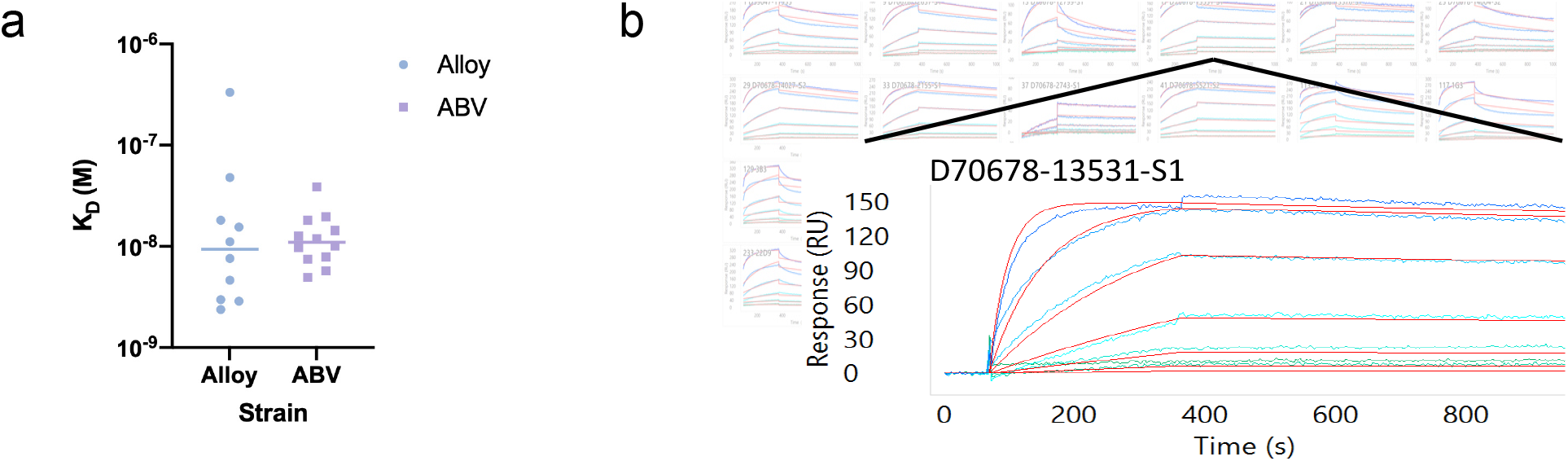
Carterra affinity analysis and example sensogram. With antibody captured on the chip surface, various concentrations of the target S1 protein were assessed for association and dissociation rates to calculate (a) final K_D_ values. (b) Array view of sensograms for each clone in the background with an example sensogram highlighted in the foreground. Each colored line indicates a distinct analyte concentration with red lines representing the curve fit analysis for rate constant calculations.

Additionally, candidates were assayed on the Carterra LSA for the ability to block the S1:ACE2 binding interaction (Table 1 and Figure 3). ACE2 receptor blocking activity was interrogated by forming an antibody:S1 protein complex on the chip surface in a sequential format. Following complex formation, ACE2 was introduced as an analyte at 100nM (Figure 3) and the percent increase in RU value as a result of ACE2 binding was quantified using the RU signal from the antibody:S1 complex formation as the baseline.

**Figure 3.**
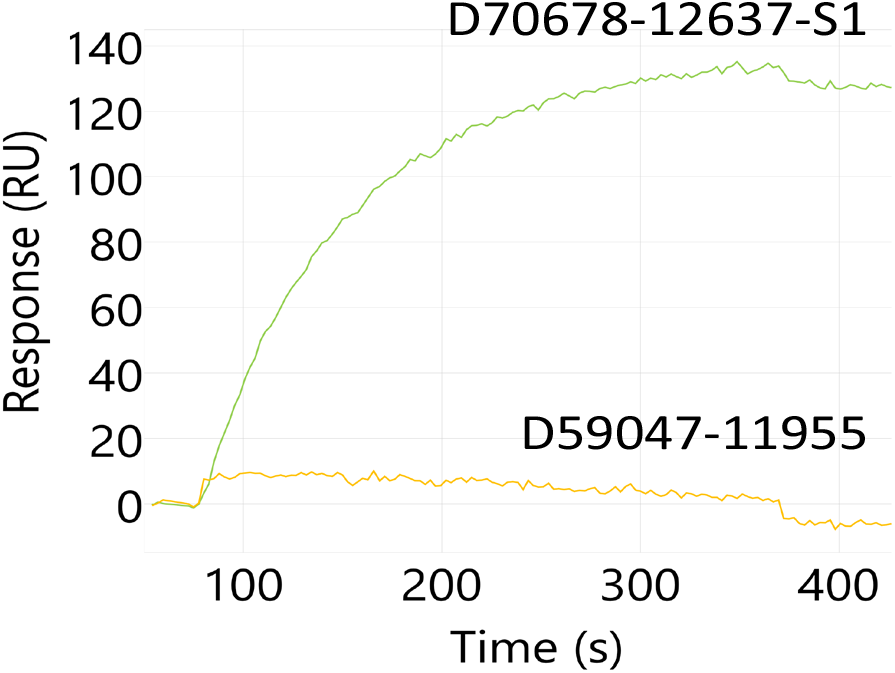
Example Carterra sensogram for antibody blocking of the S1:ACE2 interaction. Antibody:S1 protein complex was captured on the chip surface and ACE2 protein binding was assessed. A non-blocking candidate, D70678-12637-S1, complexed with S1 protein does not inhibit interaction with ACE2 (green). Conversely, a blocking candidate, D59047-11955, prevents ACE2 binding when S1 is complexed with the antibody (yellow).

The average of triplicate measurements is reported in Table 1. Non-blocking candidates resulted in a 50.8% ± 15.3% increase in signal from ACE2 binding, while blocking candidates completely prevented ACE2 binding (−1.96% ± 4.40% change in signal). One candidate, 1E5, demonstrated intermediate blocking characteristics with a 16.6% increase in signal upon ACE2 binding, perhaps indicative of a weak or partial blocking profile.

Lastly, candidate antibodies were assayed in a classical binning competition format (Figure 4) to identify the relative binding epitopes on the S1 protein. Epitope binning was performed on the Carterra LSA in a sequential format and binding of each antibody combination was assessed simultaneously. Non-competitive binding pairs and competitive binding pairs are highlighted by green or red squares, respectively, in Figure 4a (self interactions shown in dark red squares). The data is also presented as a community network plot in Figure 4b to better visualize the relative binding epitopes among the candidates assessed.

**Figure 4.**
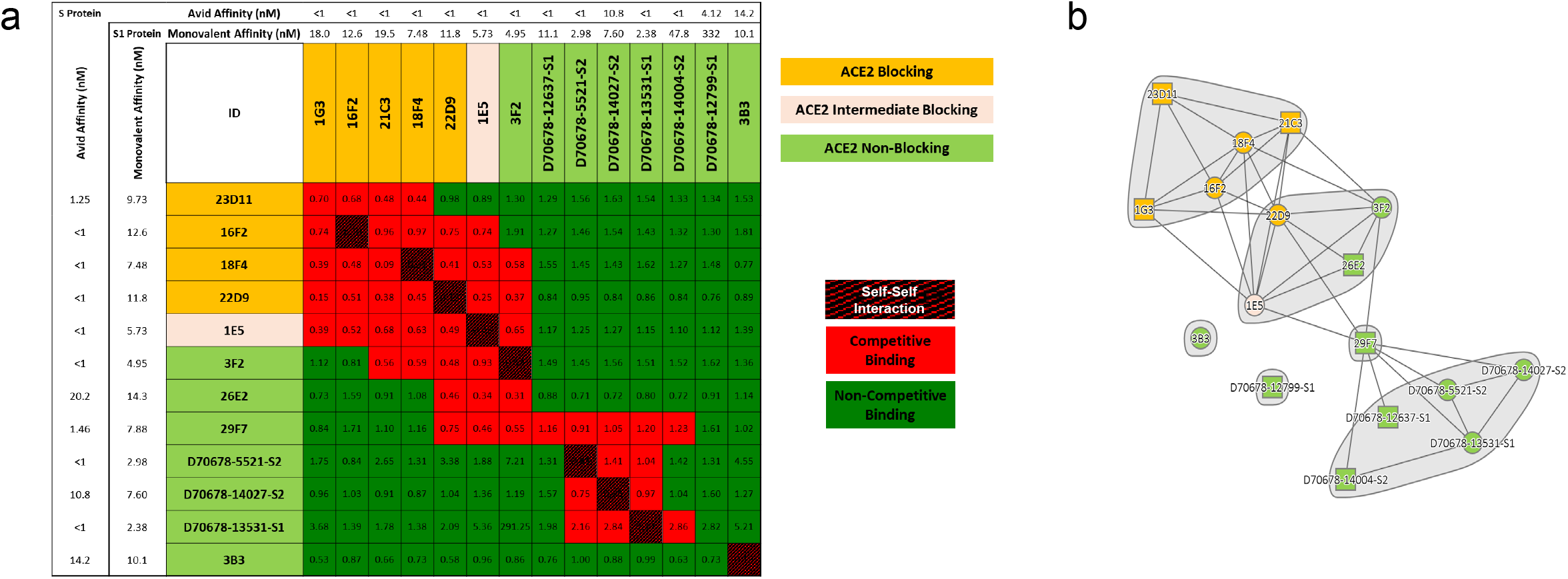
Select antibodies were characterized for competitive binding to the S1 protein to elucidate epitopic coverage. (a) All antibodies capable of binding S1 protein and preventing the S1:ACE2 interaction (ID highlighted in yellow) focused on a similar epitope (competitive binding indicated by red squares in grid). Antibodies that did not function as ACE2 blocking candidates (ID highlighted in light green) were distributed across two distinct core epitopes, with some antibodies binding at the interface of these epitopes, and an additional two clones appeared to bind distinct epitopes. Interestingly, an intermediate S1:ACE2 blocking candidate (1E5; ID highlighted in beige) bound at the interface between the ACE2 blocking epitope and a separate non-blocking epitope, thus supporting the partial blocking observation. No direct correlation was observed between affinity and binding epitope when assessed in either the monovalent binding format to the monomeric S1 protein or avid binding to the trimeric S protein. (b) A community network plot illustrates the bin clustering and distinct binding regions for each group of candidates.

Sequence information for each candidate is presented in Table 2, including full paired heavy and light chain variable regions along with V-region and mutation rate analysis. Table 3 highlights common *in silico* liability assessments for each candidate based on published motifs^10,11^ for antibodies.

**Table 2.**
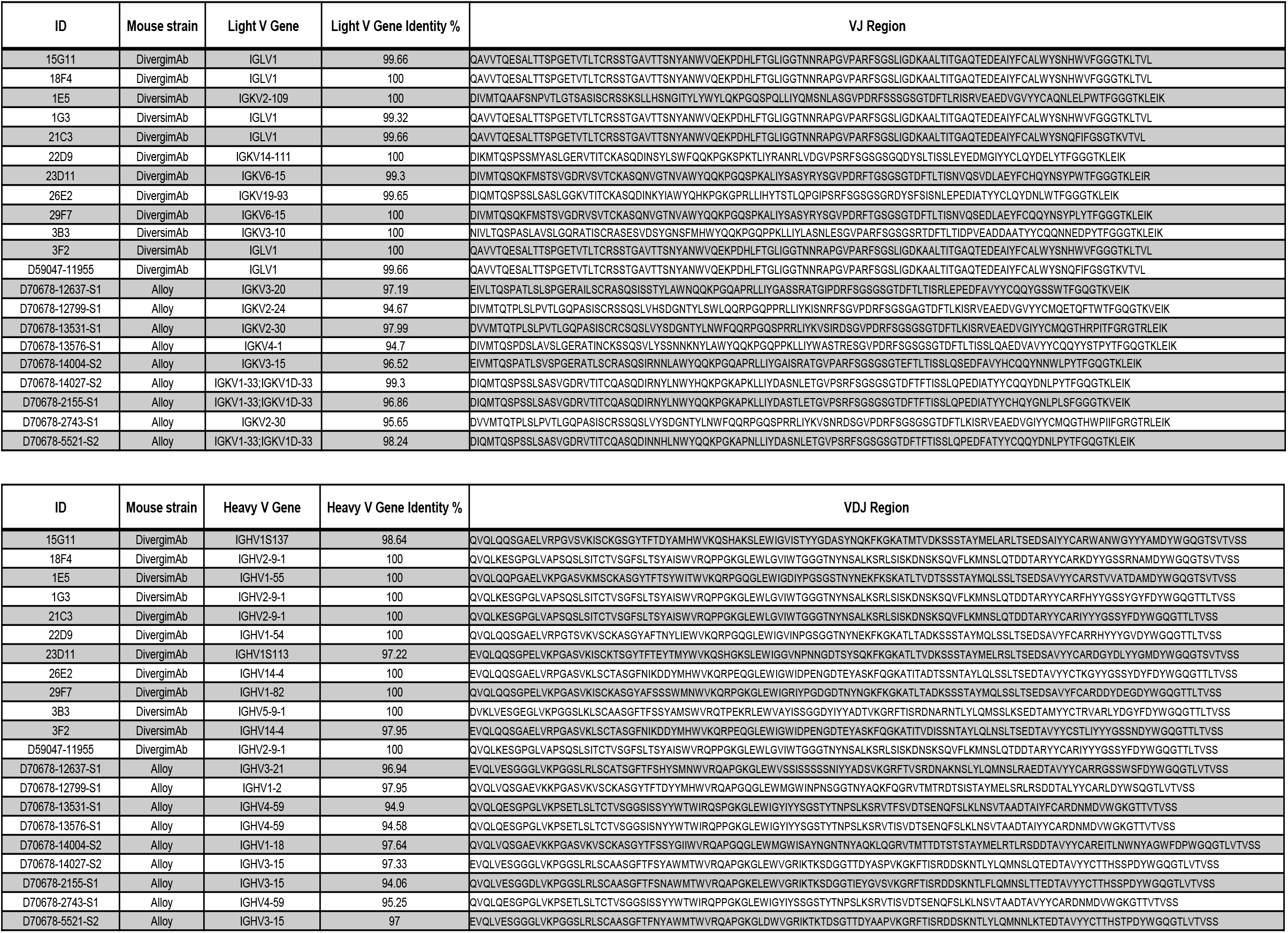
Heavy and light chain family and full sequence information for select characterized candidate antibodies.

**Table 3.** *In silico* sequence analysis of candidate antibodies for common liability motifs.

| ID | Mouse strain | Liability (High) | Liability (Medium) | Liability (Low) |
| --- | --- | --- | --- | --- |
| 15G11 | DivergimAb | DS (Isomerization, Heavy FR3); M (Oxidation, Heavy CDR3) | TS (Cleavage, Heavy FR3, Heavy FR4); TS (Cleavage, Light CDR1, Light FR1); NH (Deamidation, Light CDR3) | SN (Deamidation, Light CDR1, Light CDR3); TN (Deamidation, Light CDR2) |
| 18F4 | DivergimAb | DP (Cleavage,IGHG2B); NA (Deamidation, Heavy CDR3); NS (Deamidation, 3 * Heavy FR3); M (Oxidation, Heavy CDR3) | TS (Cleavage, Heavy CDR1, Heavy FR4); TS (Cleavage, Light CDR1, Light FR1); NH (Deamidation, Light CDR3) | TN (Deamidation,IGHV2-9-1); SN (Deamidation, Light CDR1, Light CDR3); TN (Deamidation, Light CDR2) |
| 1E5 | DiversimAb | DS (Isomerization, Heavy FR3); M (Oxidation, Heavy CDR3); NG (Deamidation, Light CDR1); M (Oxidation, Light CDR2) | TS (Cleavage, Heavy CDR1, 2 * Heavy FR3, Heavy FR4); TS (Cleavage, Light FR1); NP (Hydrolysis, Light FR1) | TN (Deamidation,IGHV1-55); SN (Deamidation, IGKV2-109, Light CDR1, Light FR1) |
| 1G3 | DiversimAb | NS (Deamidation, 3 * Heavy FR3,IGHG1) | TS (Cleavage, Heavy CDR1); TS (Cleavage, Light CDR1, Light FR1); NH (Deamidation, Light CDR3) | TN (Deamidation,IGHG1,IGHV2-9-1); SN (Deamidation, Light CDR1, Light CDR3); TN (Deamidation, Light CDR2) |
| 21C3 | DivergimAb | NS (Deamidation, 3 * Heavy FR3) | TS (Cleavage, Heavy CDR1); TS (Cleavage, Light CDR1, Light FR1) | TN (Deamidation,IGHV2-9-1); SN (Deamidation, Light CDR1, Light CDR3); TN (Deamidation, Light CDR2) |
| 22D9 | DivergimAb | DS (Isomerization, Heavy FR3); NS (Deamidation, Light CDR1); DG (Isomerization, Light FR3) | TS (Cleavage, Heavy FR1, Heavy FR3); NH (Deamidation,IGHG1); NP (Hydrolysis, Heavy CDR2) | TN (Deamidation, Heavy CDR1,IGHV1-54) |
| 23D11 | DivergimAb | NG (Deamidation, Heavy CDR2); DG (Isomerization, Heavy CDR3); DS (Isomerization, Heavy FR3); M (Oxidation, Heavy CDR3); NS (Deamidation, Light CDR3) | TS (Cleavage, Heavy FR1, Heavy FR3, Heavy FR4,IGHV1S113); NP (Hydrolysis, Heavy CDR2); TS (Cleavage, Light FR1) | SN (Deamidation, Light FR3); TN (Deamidation, Light CDR1) |
| 26E2 | DivergimAb | DP (Cleavage, Heavy CDR2); NG (Deamidation, Heavy CDR2); DS (Isomerization,IGHG2A;IGHG2C) | TS (Cleavage, 2 * Heavy FR3,IGHG2A;IGHG2C) | SN (Deamidation, Heavy FR3); SN (Deamidation, Light FR3) |
| 29F7 | DivergimAb | NG (Deamidation, Heavy FR3); DG (Isomerization, Heavy CDR2); DS (Isomerization, Heavy FR3,IGHG2C); NS (Deamidation, Light CDR3) | TS (Cleavage, Heavy FR3); TS (Cleavage, Light FR1) | TN (Deamidation,IGHV1-82); SN (Deamidation, Light FR3); TN (Deamidation, Light CDR1) |
| 3B3 | DiversimAb | NA (Deamidation, Heavy FR3); DG (Isomerization, Heavy CDR3); DP (Cleavage, Light CDR3, Light FR3); NS (Deamidation, Light CDR1); DS (Isomerization, Light CDR1) |  | SN (Deamidation, IGKV3-10) |
| 3F2 | DiversimAb | DP (Cleavage, Heavy CDR2); NG (Deamidation, Heavy CDR2); NS (Deamidation, Heavy FR3); | TS (Cleavage, Heavy FR3); TS (Cleavage, Light CDR1, Light FR1); NH (Deamidation, Light CDR3) | SN (Deamidation, Heavy CDR3, Heavy FR3); SN (Deamidation, Light CDR1, Light CDR3); TN (Deamidation, Light CDR2) |
| D59047-11955 | DivergimAb | NS (Deamidation, 3 * Heavy FR3) | TS (Cleavage, Heavy CDR1); TS (Cleavage, Light CDR1, Light FR1) | TN (Deamidation,IGHV2-9-1); SN (Deamidation, Light CDR1, Light CDR3); TN (Deamidation, Light CDR2) |
| D70678-12637-S1 | Alloy | NA (Deamidation, Heavy FR3); NS (Deamidation, 2 * Heavy FR3); DS (Isomerization, Heavy FR3) | TS (Cleavage, Heavy FR1) | KN (Deamidation, Heavy FR3); SN (Deamidation, Heavy CDR2) |
| D70678-12799-S1 | Alloy | NS (Deamidation, Heavy CDR2); DG (Isomerization, Light CDR1); M (Oxidation, Light CDR3) | TS (Cleavage, Heavy FR3); NP (Hydrolysis, Heavy CDR2) | TN (Deamidation,IGHV1-2); SN (Deamidation, IGKV2-24) |
| D70678-13531-S1 | Alloy | NS (Deamidation, Heavy FR3); M (Oxidation, Heavy CDR3); 2 C (Extra Cysteine, Light FR1); DG (Isomerization, Light CDR1); DS (Isomerization, Light FR3); M (Oxidation, Light CDR3) | TS (Cleavage, Heavy FR3); NP (Hydrolysis, Heavy FR3) | TN (Deamidation, Heavy FR3) |
| D70678-13576-S1 | Alloy | NS (Deamidation, Heavy FR3); M (Oxidation, Heavy CDR3); 2 C (Extra Cysteine, Light FR1); DG (Isomerization, Light CDR1); DS (Isomerization, Light FR3); M (Oxidation, Light CDR3); DS (Isomerization, Light FR1) | TS (Cleavage, Heavy FR3); NP (Hydrolysis, Heavy FR3) | SN (Deamidation, Heavy CDR1); TN (Deamidation, Heavy FR3); KN (Deamidation, Light CDR1); SN (Deamidation, Light CDR1) |
| D70678-14004-S2 | Alloy | DP (Cleavage, Heavy CDR3); NG (Deamidation, Heavy CDR2); | TS (Cleavage, 2 * Heavy FR3) | TN (Deamidation,IGHV1-18) |
| D70678-14027-S2 | Alloy | NS (Deamidation, Heavy FR3); DG (Isomerization, Heavy CDR2); DS (Isomerization, Heavy FR3); |  | KN (Deamidation, Heavy FR3); SN (Deamidation, IGKV1-33;IGKV1D-33) |
| D70678-2155-S1 | Alloy | NA (Deamidation, Heavy CDR1); NS (Deamidation, Heavy FR3); DG (Isomerization, Heavy CDR2); DS (Isomerization, Heavy FR3); |  | KN (Deamidation, Heavy FR3); SN (Deamidation, Heavy CDR1) |
| D70678-2743-S1 | Alloy | NS (Deamidation, Heavy FR3); M (Oxidation, Heavy CDR3); DG (Isomerization, Light CDR1); DS (Isomerization, Light FR3); M (Oxidation, Light CDR3) | TS (Cleavage, Heavy FR3); NP (Hydrolysis, Heavy FR3) | TN (Deamidation, Heavy FR3); SN (Deamidation, IGKV2-30) |
| D70678-5521-S2 | Alloy | DS (Isomerization, Heavy CDR2, Heavy FR3) | NH (Deamidation, Light CDR1) | KN (Deamidation, Heavy FR3); SN (Deamidation, IGKV1-33;IGKV1D-33) |

## Discussion

The COVID-19 pandemic highlighted the importance of rapid discovery of antiviral drugs, both prophylactic and therapeutic, as critical to containing the spread of the virus.^12^ In response to the COVID-19 outbreak, the scientific community collectively answered the public health call to action with extraordinary momentum, leveraging both new and traditional technologies with a strong emphasis on speed.^13^ Continuing to build on COVID-19 research critical to public health while highlighting various methods that can be deployed for antibody discovery, this report outlines the use of a number of distinct workflows that enabled the accelerated identification of antiviral antibodies, some with promising therapeutic potential. The sequence information and corresponding characterization data for a panel of 21 antibody candidates is reported for unrestricted use.

We highlight the use of both DiversimAb and Alloy GK mice in accelerated hybridoma-based and single B cell screening platforms. To enable rapid generation of monoclonal antibodies out of the DiversimAb platform, mice were immunized on a 16-day accelerated schedule followed by hybridoma generation. Concurrently, Alloy mice were immunized on a 5-week schedule and subsequently screened on the Beacon in a single day followed by a sequence recovery and analysis process spanning less than one week. Despite the divergent immunization and screening workflows, both campaigns yielded sequences in similar timeframes to enable simultaneous downstream characterization requiring three total days. In sum, both campaigns required fewer than three months from immunization start to fully-characterized, purified antibody. Of course, alternative workflows are possible to further tighten the timeline for future campaigns. For example, characterization data was acquired using both purified and crude antibody samples to validate either source on the Carterra LSA for kinetic, neutralization and binning data. With equivalent results, the time and resources required for purification can be incorporated further downstream to expedite the early discovery timeline. In addition, Alloy mice can be immunized on a more accelerated timeline.

The resulting data set presented here is comprised of a diverse panel of candidate antibodies spanning multiple epitopes with high affinity and both receptor blocking and non-blocking activity. Interestingly, all candidate antibodies identified from the Alloy mice fell within a similar non-blocking bin, likely indicative of epitope immunodominance,^14^ which has been reported as a common result from COVID-19 infection.^15,16^ However, the extended immunization strategy employed for the Alloy mice resulted in high affinity antibodies, including the top four highest affinity candidates presented here. The balance between diversity and affinity is a challenge for *in vivo* immunization models: with continued boosting, clonal expansion of the highest affinity germline B cells can often result in a limited overall diversity.^17^ Alternative immunization workflows involving rapid schedules and/or immunogen manipulation are effective risk mitigation strategies to circumvent these challenges.^18^ In this case, the DiversimAb mice were immunized following an accelerated strategy to maximize epitopic diversity, and although the overall average affinity of these candidates was lower as a result, a subset of high affinity blocking clones were discovered. It is also important to note that recent studies have identified non-blocking neutralizing epitopes,^19^ indicating further testing of non-blocking Alloy candidates identified here could reveal efficacious potential in viral neutralizing experiments.

A review of the lead candidate sequences further underscores the importance of risk mitigation in antibody discovery through the use of multiple strains of mice for added diversity. It is well documented that genetic backgrounds heavily influence the B cell repertoire diversity.^20,21^ Therefore, in cases where rapid discovery is imperative, starting with multiple strains can improve the diversity in gene usage for V(D)J recombination. In the case of the Abveris DiversimAb mice, two separate background strains were leveraged (DiversimAb vs. DivergimAb) to further increase the output sequence diversity. Interestingly, many of the candidates from these mice contained near germline V-regions - likely a result of the accelerated immunization approach employed for rapid discovery. Regardless, binding affinity is still maintained at the nanomolar level, which could be due to the unique and diverse CDR3 regions. The presence of germline sequences from DiversimAb mice using this immunization approach provides an opportunity for streamlined lead optimization via humanization and affinity maturation without the need to assess numerous V region backmutation permutations.

Traditional methods of *in vivo* antibody drug discovery suffer from timeline disadvantages associated with immunization, humanization, and downstream lead optimization (if required).^22^ However, with the recent development of innovative new technologies to accelerate the upstream drug discovery process highlighted in this report, *in vivo* antibody discovery is now a viable option for an accelerated response to novel viral threats. Humanized mouse strains can be leveraged synergistically with genetically engineered mice designed to increase epitopic diversity (DiversimAb) to provide lead and backup antibody drug candidates of desired therapeutic efficacy. When combined with efficient high-throughput downstream antibody capture and characterization tools, it becomes possible to go from immunization to sequence in as few as 29 days.

## Acknowledgements

We would like to thank Alloy Therapeutics for supplying the GK mice, Carterra for assistance and use of the LSA and Gary Ng of Abveris for preparation of the manuscript.

## Author Contribution

TE.M. and C.A.S. conceived of and designed experiments, performed data analysis, and authored the manuscript. R.A., J.B., A.S.B., N.K.D., N.T.D., A.M.D., C.E., J.G., B.G., K.S.P., K.R., and J.S. designed and executed experiments, performed data analysis and reporting, and participated in manuscript review.

## Supplementary Methods

### Mouse immunization, B-cell enrichment and hybridoma generation

Mouse immunizations were performed at Abveris (Canton, MA) in accordance with all IACUC protocols. Eight- to twelve-week-old DiversimAb or DivergimAb mice received immunizations following a RIMMS schedule at subcutaneous sites using a custom adjuvant formulation reconstituted in an oil-in-water emulsion over the course of 18 days. Twelve-week-old Alloy GK+ mice (Alloy Therapeutics) received immunizations following a weekly schedule at intraperitoneal and subcutaneous sites using Freund’s adjuvant over the course of 5 weeks. S1 protein for immunization (Acro) was used at a dosage ranging from 10-100 μg per injection. Titer tests were performed via indirect ELISA to S1 and S proteins (Acro) to select mice with the highest antigen-specific titer. Mice received a final boost 1 to 3 days before harvest of lymph nodes and spleens. For single B cell workflows, plasma B cells were isolated from a B cell enriched population using a magnetic plasma cell isolation kit. Isolated B cells were resuspended at the recommended density for Beacon (Berkeley Lights) import and stained for targeted penning. Hybridoma generation followed standard electrofusion techniques.^23^ Briefly, single cell suspensions from tissue isolations were combined with SP2/0 myeloma cells and fused via electrofusion (BTX). Resulting hybridoma clones were distributed into 96-well plates and expanded over 10-14 days with HAT (Sigma-Aldrich) selection prior to screening.

### Single B cell screening

#### Cell loading

Plasma B cells were imported onto 14k OptoSelect chips following a modified Opto Plasma B Discovery workflow. Briefly, through a customized Targeted Pen Selection (TPS) penning algorithm, live plasma B cells were penned via OEP™ technology as single cells over multiple rounds of imports, while pens containing multiple cells were unpenned. Cells were penned in a custom medium formulation containing Cell MAb Medium Quantum Yield (ThermoFisher Scientific) supplemented with FBS, growth factors and Loading Reagent (Berkeley Lights). Assays and culture were performed under the same media conditions, but in the absence of Loading Reagent.

#### Bead-based screening assays

Prior to Beacon workflows, recombinant S1 (Acro) and ACE2 (Acro) proteins were labeled at primary amine sites with AF488 or AF647 using Alex Fluor 488 TFP Ester or Alexa Fluor 647 NHS Ester, respectively (ThermoFisher Scientific). For screening and ligand blocking assays, chips containing penned plasma B cells were loaded with a single or multiplexed assay mixture containing anti-mouse IgG coated beads (Spherotech) and fluorescently labeled target protein(s) at optimized concentrations (200nM or 500nM for S1 and ACE2 proteins, respectively) predetermined in bio-layer interferometry optimization assays. Standard AbDisc2.0 Beacon assay import conditions were used and chips were imaged over the course of 30 to 60 minutes in 2 minute intervals between images. Over the course of the assays, secreted antibody from plasma B cells diffused from the NanoPen chambers into the channel where they bound the beads and concentrated fluorescently labeled protein(s) or secondary antibodies, forming fluorescent halos in the channels adjacent to the pens containing antigen-specific plasma cells. The chips were imaged using a FITC and/or Cy5 filter cube. For sequential assays, the bead assay mixture was flushed from the chip before loading a second assay mixture. All assays were automatically scored and confirmed with human verification.

#### Sequence recovery and antibody expression

Cells of interest identified via Beacon screening were selected for automated single cell export into 96-well PCR plates containing lysis buffer. Selected hybridoma clones were pelleted and washed in PBS prior to lysis. Paired heavy and light chain sequences were amplified using genespecific primers for cDNA production and two successive rounds of nested PCR. Nucleotide sequence from amplicons was determined by Sanger sequencing in reverse orientation and data was analyzed in Geneious Biologics (Biomatters) for antibody annotation using a customized Single Clone Antibody Analysis pipeline. Annotations for closest germline V(D)J genes, degree of somatic hypermutation and potential sequence liabilities were extracted. For transient recombinant expression of single cell workflow candidates, paired heavy and light chain sequences were codon optimized, synthesized (ThermoFisher Scientific), cloned into separate mammalian expression vectors, sequenced to confirm identity and correctness, then co-transfected into Expi293™ cells (ThermoFisher Scientific) for supernatant harvest 5 days post-transfection. Candidates from hybridoma clones were scaled to sufficient volume in media containing low IgG FBS (Corning) and hybridoma viability was monitored to indicate optimal supernatant harvest dates. Antibody presence in saturated supernatant from recombinant expression or hybridoma cultures was confirmed by bio-layer interferometry (ForteBio Sartorius), purified by ProteinA/G (Cytiva), and quantitated by UV absorption spectrophotometry.

#### Primary screening by ELISA and bio-layer interferometry

Supernatant from recombinant expression or hybridoma cultures was screened by indirect ELISA to S protein and counter screened against an irrelevant Avi- and HIS-tagged protein (Abveris). Target proteins were coated on high absorption plates (Corning) at 1 μg/ml, washed and incubated with supernatant samples at room temperature. Mouse IgG specific to the coated proteins was detected with HRP-conjugated anti-mouse IgG Fc-specific detection antibody (Jackson Immunoresearch). Supernatant from positive candidates based on ELISA screening was subsequently assayed on the Octet RED96e (ForteBio Sartorius) using AMC biosensors. Immobilization of mouse IgG onto AMC biosensor surfaces was performed for 120 seconds, followed by association to 100nM of trimeric S protein analyte for 300 seconds and dissociation in buffer for 600 seconds. All steps were performed at 25°C and 1000 rpm. Data was analyzed using Analysis HT software v11.0 with double referencing to irrelevant polyclonal mIgG (Jackson Immunoresearch) and irrelevant Avi- and HIS-tagged protein (Abveris). Full curve fits were assessed with Savitzky-Golay Filtering using a 1:1 model.

#### High-content screening by surface plasmon resonance

Candidates of interest based on primary screening results were assessed by high-throughput surface plasmon resonance (SPR) on a Carterra^®^ LSA™ platform. Supernatant from recombinant expression and hybridoma cultures, as well as corresponding purified antibodies, were measured in triplicate via a capture format on CMD200M (Carterra) or HC30M (Carterra) sensor chip. In this format, 50 μg/ml of goat anti-mouse Fc antibody (Jackson Immunoresearch) in NaOAc pH 4.5 buffer was coupled to the chip surface via standard EDC-NHS activation. Experimental and control antibodies were subsequently printed in an array format on the capture lawn for 30 minutes diluted to 10 μg/ml for purified antibodies or 1:1 dilution for supernatant samples in HBSTE (Carterra). Captured antibodies were subsquently crosslinked with 870 μM BS3 (ThermoFisher Scientific) to enable regenerative conditions in subsequent steps without the requirement to re-array experimental and control antibodies.Both non-regenerative (single cycle) and regenerative (multicycle) kinetics analyses were performed using monomeric S1 protein analyte with six sequential injections at different concentrations ranging from 0.46nM to 500nM in HBSTE + 0.1% BSA running buffer. Analyte was injected for 5 minutes to measure association followed by 15 minutes of buffer injection to assess dissociation. For multi-cycle kinetics, standard regeneration conditions were used with alternating injections of 10mM glycine pH2.5 and running buffer.

To assess ACE2 blocking profiles, a single injection of 100nM ACE2 in running buffer was performed following S1 binding at 500nM to the captured antibodies on the chip surface. Using the RU value of antibody:S1 complex as a baseline, the increase in RU response due to ACE2 binding was calculated as a percentage of the baseline. Kinetic and blocking data were analyzed using Kinetics software v1.6 and association and dissociation rate curve fits were calculated using a 1:1 Langmuir binding model.

An epitope competition assay was performed in the classical sandwich format whereby the captured and crosslinked array of experimental and control antibodies was sequentially exposed to the following series of analyte injections: 200 μg/ml polyclonal mIgG for 5 minutes to block additional surface sites, 100nM S1 protein in running buffer for 5 minutes, 10 μg/ml (purified) or 1:1 dilution (supernatant) in running buffer of an individual experimental or control antibody. Each injection series was preceded by and ended with a buffer injection. Standard regeneration conditions were used between each full cycle. Analyte injection series were repeated until all experimental and control antibodies were assessed as analytes. Crude supernatant and purified antibodies were assessed in bi-directional format as both ligands and analytes. Binning data was analyzed using Epitope software v1.6 with manual verification to generate clustered heat map and community network diagrams. Competition for S1 binding was based on normalized signal to the S1 binding step for each interaction and self-self-competition was used as a baseline for determining positive binding. Samples with sufficient S1 protein capture and adequate regeneration profiles were reported with bi-directional data, otherwise a unidirectional interaction (analyte only) was reported.

